# Comparative Pore Structure and Dynamics for Bacterial Microcompartment Shell Protein Assemblies in Sheets or Shells

**DOI:** 10.1101/2024.03.12.584231

**Authors:** Saad Raza, Daipayan Sarkar, Leanne Jade G. Chan, Joshua Mae, Markus Sutter, Christopher J. Petzold, Cheryl A. Kerfeld, Corie Y. Ralston, Sayan Gupta, Josh V. Vermaas

## Abstract

Bacterial microcompartments (BMCs) are protein-bound organelles found in some bacteria which encapsulate enzymes for enhanced catalytic activity. These compartments spatially sequester enzymes within semi-permeable shell proteins, analogous to many membrane-bound organelles. The shell proteins assemble into multimeric tiles; hexamers, trimers, and pentamers, and these tiles self-assemble into larger assemblies with icosahedral symmetry. While icosahedral shells are the predominant form *in vivo*, the tiles can also form nanoscale cylinders or sheets. The individual multimeric tiles feature central pores that are key to regulating transport across the protein shell. Our primary interest is to quantify pore shape changes in response to alternative component morphologies at the nanoscale. We use molecular modeling tools to develop atomically detailed models for both planar sheets of tiles and curved structures representative of the complete shells found *in vivo*. Subsequently, these models were animated using classical molecular dynamics simulations. From the resulting trajectories, we analyzed overall structural stability, water accessibility to individual residues, water residence time, and pore geometry for the hexameric and trimeric protein tiles from the *Haliangium ochraceum* model BMC shell. These exhaustive analyses suggest no substantial variation in pore structure or solvent accessibility between the flat and curved shell geometries. We additionally compare our analysis to hydroxyl radical footprinting data to serve as a check against our simulation results, highlighting specific residues where water molecules are bound for a long time. Although with little variation in morphology or water interaction, we propose that the planar and capsular morphology can be used interchangeably when studying permeability through BMC pores.

## Introduction

Bacterial microcompartments (BMC) are self-assembling protein based organelles found in various bacteria.^1,2^ BMCs are thought to have evolved to facilitate catalysis for difficult or dangerous reactions. The semi-permeable BMC shell protects the bacterial cytosol from toxic effects of unstable or reactive intermediates that are present in these metabolic pathways by sequestering these intermediate products.^3^ Spatially confining these reaction pathways also increases their metabolic efficiency, in part by creating local high concentrations for enzyme substrates.^4^ Enzyme compartmentalization increases the rate of catabolism which increases fitness.^5^ BMC shells also serve to protect encapsulated enzymes from deleterious metabolites, such as O_2_ for oxygen sensitive enzymes.^6,7^ For all of these reasons, BMC shell are emerging as engineering platforms for abiotic and biotic catalysis. ^8–11^

A crucial limitation for engineering new catalytic pathways into BMC shell are whether reactants and products permeate across the shell. Natural BMCs are permeable to a wide range of metabolites, such as bicarbonate and Calvin-Benson-Bassham cycle intermediates to facilitate carbon fixation in cyanobacterial carboxysomes,^12^ or reactants and products for propanediol,^13^ ethanomine,^14^ frucose or rhamnose^15^ synthesis. It is thought that the permeation occurs through the central pores within the individual tiles identified from molecular structures,^16,17^ corroborated by molecular simulation for metabolites through these pores.^18–20^

Molecular simulation can model the permeation for any metabolite across these pores explicitly. A common approach is to use isolated proteins tiles in solution as a model for the BMC shell, effectively measuring permeability in the dilute limit. ^19,20^ More recently, intact shells have been simulated.^18^ While intact shells are a more accurate representation for the molecular nanostructure, simulating these assemblies at the atomic scale substantially raises the cost of determining molecular permeability. The increased computational cost is particularly acute when trimeric shell components are included, as the shell size increases substantially (Fig. 1). Since most of the simulation volume for these intact shells is water, an alternative intermediate system that reflects the symmetry of shell components would be ideal to better balance computational cost and accuracy.

**Figure 1:**
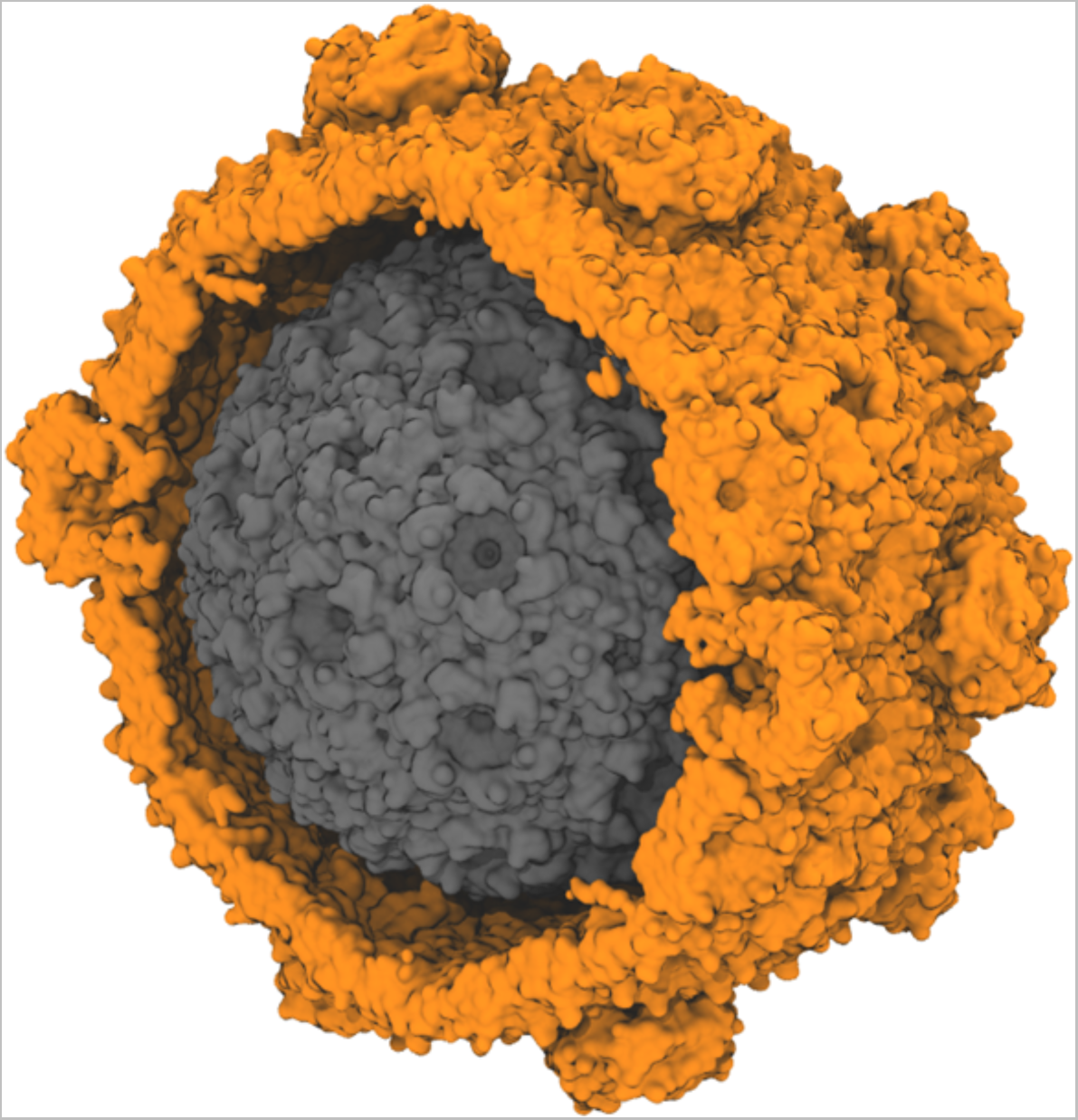
Size comparison for different BMC shell architectures made from *Haliangium ochraceum* shell proteins. The shown structures are determined by cryo-EM, and represent a minimal shell (grey, PDB:6OWG, approximately 2.4M atoms when solvated for simulation)^21^ and a full HO shell (orange, PDB:6MZX, approximately 10M atoms when solvated for simulation).^22^ The minimal shell contains only of hexameric and pentameric units while a typical full shell also includes trimeric proteins, some of which form double-stacks in the shell. The diameter of the minimal model BMC is around 22 nm, while the full shell diameter is approximately 40 nm.

To explore this idea, we leverage the fact that BMC shell proteins do not always form shells. Alternative morphologies such as sheets of hexamers have been experimentally characterized and cylinders have been reported *in vitro* after modifying the ratio of shell components for the *Haliangium ochraceum* (HO) shell. In this study, we compare the dynamics and structure of a small periodic BMC sheet to an equivalent shell fragment. Crucially, we find that the dynamics and pore diameter for a planar or shell-structures to be similar, indicating that the permeability for the developed sheet system is similar to the permeability that would be expected *in vivo*.

## Methods

### Structure Preparation

Fundamentally, there are two different models prepared in this study, a curved facet from a larger BMC shell, and a planar arrangement of the same proteins. The curved facet (labeled as shell in Fig. 2) starts from existing structures determined from cryo-EM, specifically the 6N07 structure that features a single stacked BMC trimer surrounded by BMC hexamers.^22^ This starting structure is prepared in VMD^23^ using the solvate and autoionize plugins to create a 234 x 203 x 234Å system suitable for further simulation.

**Figure 2:**
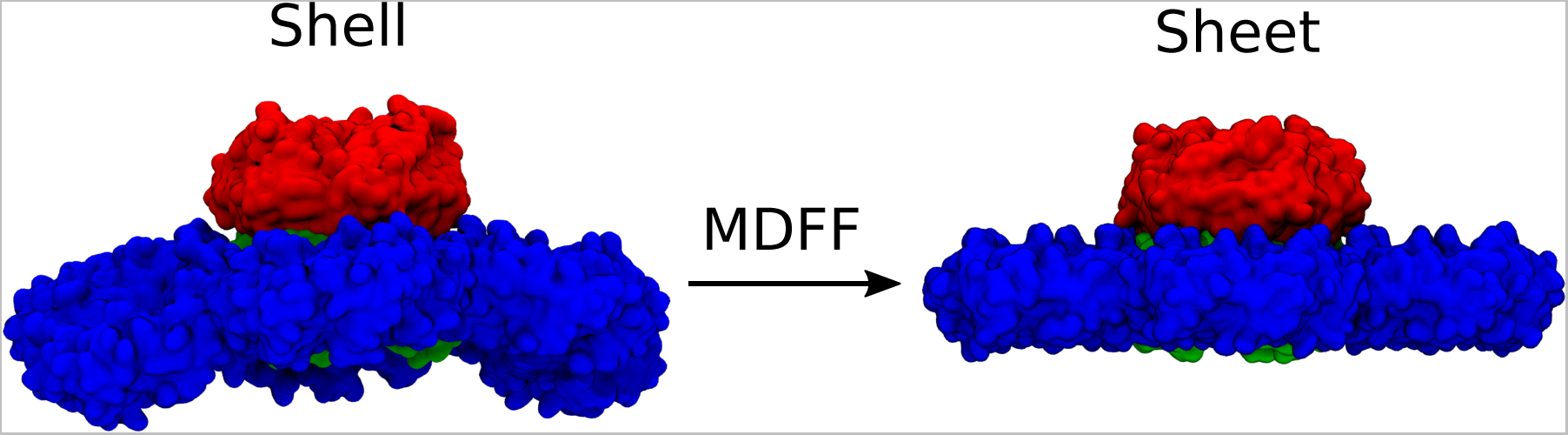
Using MDFF methods, we convert the shell-like curved facet of BMC (left) from PDB ID:6N07 to a sheet-like conformation (right).Hexamer tiles are in blue, and the stacked trimers, are a dimer of two trimers (one red, one green), resulting in the red trimer protruding from the plane of the shell facet, in which the green trimer is embedded.

To create a planar sheet-like structure, we use molecular dynamics flexible fitting (MDFF)^24,24,25^ to flatten the initial structure (Fig. 2). Since MDFF requires an electron density, either real or synthetic, as the target, we generated a nominally flat target conformation for the hexamers. Starting with the curved facet, we first determine a vector normal to the trimer pore by creating vectors 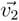 and 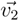 from adjacent protein pairs in the upper trimer. The cross product of these two vectors is normal to the trimer pore, and can be brought to a specific axis using the transvecinv routine in VMD.^23^ The trimer is moved to the origin, and this procedure is repeated for each hexamer individually to create a the sheet arrangement. After transformation, the synthetic target density map was generated at 4 Å resolution via the mdff plugin within VMD,^23^ using the combined atomic model of the flattened hexamers. The Tcl scripts for generating the flattened hexamers using rigid body transformation and generating the synthetic density map of the flattened system is available via Zenodo. ^26^

In order to create an effectively infinite planar sheet of BMC shell proteins, the flattened facet was arranged in the X-Y plane such that the pore normal is aligned with the Z-axis. We further reduce the system size by packing three hexamer and one trimer tile into the unit cell, such that the trimer tile is always surrounded by hexamer tiles as is the case in cryo-EM structure.^22^ The X-Y plane dimensions were set based on the distance between repeating units within the structure. Once optimized in this manner and solvated, the unit cell dimensions for the final structure in Fig. 3B are 130 x 170 x 152 Å. Fig. 3C places the repeating unit within the larger context, while Fig. S1 explicitly shows a surface representation for the repeating unit and how it tiles together.

**Figure 3:**
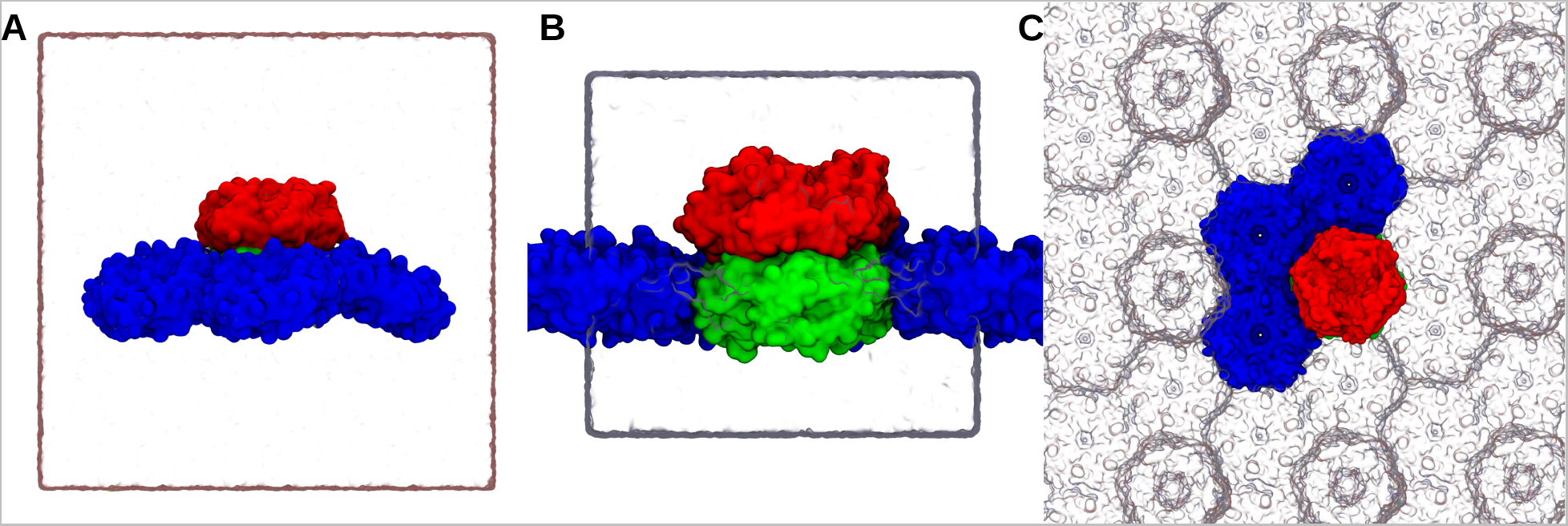
Simulation box of curved shell (A) and planar sheet conformation (B) with water box represented as the white rectangular which also represent the periodic boundary of the system. (C) Top view of sheet simulation system comprising of three hexameric units and one trimer dimer which can mimic the planar sheet conformation and keeps the system periodic in a rectangular simulation box. Periodic images of the facets are presented as glass bubble. Hexamers are represented as blue surface representation, trimer upper half as red and lower half as green surface representation. Planar conformation can be seen mimicking a lipid membrane where the hexameric units are touching the periodic trimeric subunits. Animations for these simulations systems are available as SI Animation 1&2.

### Simulation Protocol

Classical molecular dynamics simulations were carried out using the CHARMM36m protein force field^27^ in explicit TIP3 solvent^28^ using NAMD.^29^ Minimization and a brief equilibration was performed with NAMD 2.14, and 1 µs unbiased production simulations were performed using the GPU-resident integrator on NAMD 3.0a9 to maximize performance.^29^

Most simulation parameters were shared when running the shell and sheet-like structures shown in Fig. 3. Temperature was controlled using the Langevin thermostat at 298 K with 1 ps^-1^ damping. Hydrogen bonds were handled with SETTLE algorithm to enable 2 fs timesteps.^30^ Long-range non-bonded Lennard Jones (LJ) cutoff was set to 12 Å. Long-range electrostatic interactions were calculated with particle mesh Ewald (PME) grid with 1.2 Å spacing^31,32^. The switching between non-bonded interaction and electrostatic is done after 10 Å. LJ correction is applied to improve energy conservation during switching.^33^ Energy minimization of the system was initially performed using the 1000 steps of conjugate gradient in NAMD.^34^ Both systems were equilibrated for 50 ps in NPT ensemble using 5 Å margin to allow the box to adjust after any distortion from minimization prior to transitioning to the GPU-resident integrator. Simulations in the production NPT ensemble were run for 1 µs with default margin to maximize performance.

The difference between the sheet and shell structures occur in the pressure control. A Langevin barostat was used to maintain pressure at 1 atm,^35^ with the shell fragment using semi-isotropic pressure control and the sheet using anisotropic pressure control. For sheet simulations, the x- and y-dimensions are tied to expansion and contraction of the individual shell proteins. Unlike lipid bilayer systems, where the membrane plane would be uncoupled except through the barostat, the protein itself can grow and shrink by different amounts along the axes parallel to the sheet surface. Thus, shell fragment simulations used a flexible simulation box with constant ratio, while production simulations for the sheet structure applied anisotropic pressure control without constant ratio.

### X-ray Footprinting with Mass Spectrometry (XFMS)

Two samples were prepared for X-ray Footprinting with Mass Spectrometry (XFMS). The first sample was taken from intact synthetic HO shells, assembled following the steps laid out in prior literature.^36^ The second sample was the purified component HO BMC-H protein (hexamer tile), which spontaneously forms uniformly oriented sheets at high concentrations;^36,37^ this was diluted to the extent that no sheets formed.

Samples were exposed at the Advanced Light Source beamline 3.2.1 with exposures of 0, 100, 250, 500, 750, 1000 and 2000 s, using a horizontal capillary as previously described.^38^ Post-exposure, samples were digested using trypsin enzyme (Promega) overnight at 37 °C at pH 8 in 50 mM ammonium bicarbonate buffer. Liquid chromatography-mass spectrometry (LCMS) was conducted on an Agilent 6550 iFunnel Q-TOF mass spectrometer (Agilent Technologies, Santa Clara, CA) coupled to an Agilent 1290 LC system (Agilent). Peptide samples were loaded onto a Sigma-Aldrich Ascentis Peptides ES-C18 column (2.1 mm x 100 mm, 2.7 µm particle size; Sigma-Aldrich, St. Louis, MO) via an Infinity Autosampler (Agilent) with Buffer A (2% Acetonitrile, 0.1% Formic Acid) with flow rate 0.400 mL/min. Peptides were eluted into the mass spectrometer via a gradient with an initial condition of 5% buffer B (98% Acetonitrile, 0.1% Formic Acid) increasing to 90% B over 15 minutes. The data were acquired with MassHunter B.05.00 operating in Auto MS/MS mode whereby the three most intense ions (charge states 2 - 5) within m/z 300 to 1400 mass range above a threshold of 1000 counts were selected for MS/MS analysis. MS/MS spectra were collected with the quadrupole set to “Narrow” resolution and collision energy to optimize fragmentation. MS/MS spectra were scanned from m/z 100 to 1700 and were collected until 40000 total counts were collected or for a maximum accumulation time of 333 ms. Parent ions were excluded for 0.1 minutes following MS/MS acquisition.

XFMS peptide identification and analysis was performed using the Byos® (Protein Metrics Inc) integrated software platform at the Molecular Foundry as previously described.^39^ Briefly, the abundance of the identified unmodified and modified peptides at each irradiation time point area were measured from their respective extracted ion chromatogram of the mass spectrometry data collected in the precursor ion mode. The fraction unmodified for each peptide was calculated as the ratio of the integrated peak area of the unmodified peptide to the sum of integrated peak areas from the modified and unmodified peptides. The dose-response curves (fraction unmodified vs. X-ray exposure) were fitted to single exponential functions, producing a k-value (s*^−^*^1^). The ratio of k-values provided the relative change in the solvent accessibility between the sheet and shell forms.

### Solvent Accessibility Analysis

Structure files and MD simulation trajectories were visualized and analyzed using Python-enabled VMD 1.9.4a58^23^. Python enabled VMD provides an interface to apply the numpy numerical library^40^ and plotting tools like matplotlib.^41^ The stability of the system was assessed first through computing the root mean square deviation (RMSD). The solvent-accessible surface area (SASA) was computed residue-wise, accelerated by a modified analysis routine that has been committed upstream to the VMD developers. Water contacts with the hexamers and trimer was calculated using the contact function in VMD to track the number of unique water molecules within 5 Å of a given residue. Beyond computing water contacts, we also calculated the water retention time around the hexamers and trimers. The residence time was determined by tracking frame by frame if a water molecule was initially within 5 Å of a given residue, and stopping the clock when the water molecule was further than 8 Å away from the residue.

To compare directly with experimental observations based on hydroxyl radical footprinting data, we subdivided the shell fragment hexamers based on their water accessibility. For the purposes of comparing with intact shells, we evaluate hexamer monomers within the shell fragment system that are interfacing directly with the trimer tile (red in Fig. S2). When comparing to dilute hexameric tiles in solution, we compare with hexamer monomers in the shell fragment system that are solvent exposed (blue in Fig. S2). This facilitates direct comparison with the companion experiment.

### Pore Analysis

The primary analysis of interest is determining the pore size within a BMC protein tile. Borrowing from membrane protein studies, we used the HOLE program^42^ to determine pore radius along the channel formed at the center of BMC hexamer and trimers tiles. The HOLE algorithm works by finding a maximum sphere fitting inside the cavities of protein along the z axis of protein. The HOLE program was written to analyze a single conformation, and so for full trajectory analysis an additional wrapper is required. While other tools such as MDAnalysis have such wrappers already built-in,^43,44^ optional parameters were essential to guiding HOLE along the pore of interest. In this vein, we wrote HoleHelper, a Tcl plugin to VMD that facilitates using HOLE for our specific systems with VMD atomselection language. HoleHelper is available on github for public download and use (https://github.com/joshuamae/HoleHelper).^45^

## Results

Molecular simulation provides a unique perspective to address specific mechanical and structural questions at the nanoscale, and has been called a ”computational microscope”.^46,47^ Turning this microscope to BMC shell protein assemblies, the key question is whether the pores respond at all to the environment, as known for the opening and closing mechanism for mechanosensitive channels.^48^ By also checking for stability and comparing our structures to experimental observables, we are confident that the BMC shell components are closer in nature to aquaporins, and do not depend on external pressure to open or close.

### Structural Stability Considerations

Prior to any pore geometry comparison, we use the root mean square deviation (RMSD) over time to assess for general protein stability within our simulation environment. Since the resolution for the original cryo-EM structure is 3.6 Å,^22^ we anticipate a RMSD similar in magnitude to this resolution, as this relationship has been noted previously for membrane proteins.^49^ That is indeed what we see (Fig. 4), with extended simulation only yielding RMSDs that occasionally exceed the solved structure resolution. The shell fragment routinely has lower RMSD than the sheet, suggesting that there are subtle structural changes that have occurred, as the reference structure for each tile is identical between both states. However, since the RMSD change is so small, and largely confined to the hexamers, the overall secondary structure is consistent between states.

**Figure 4:**
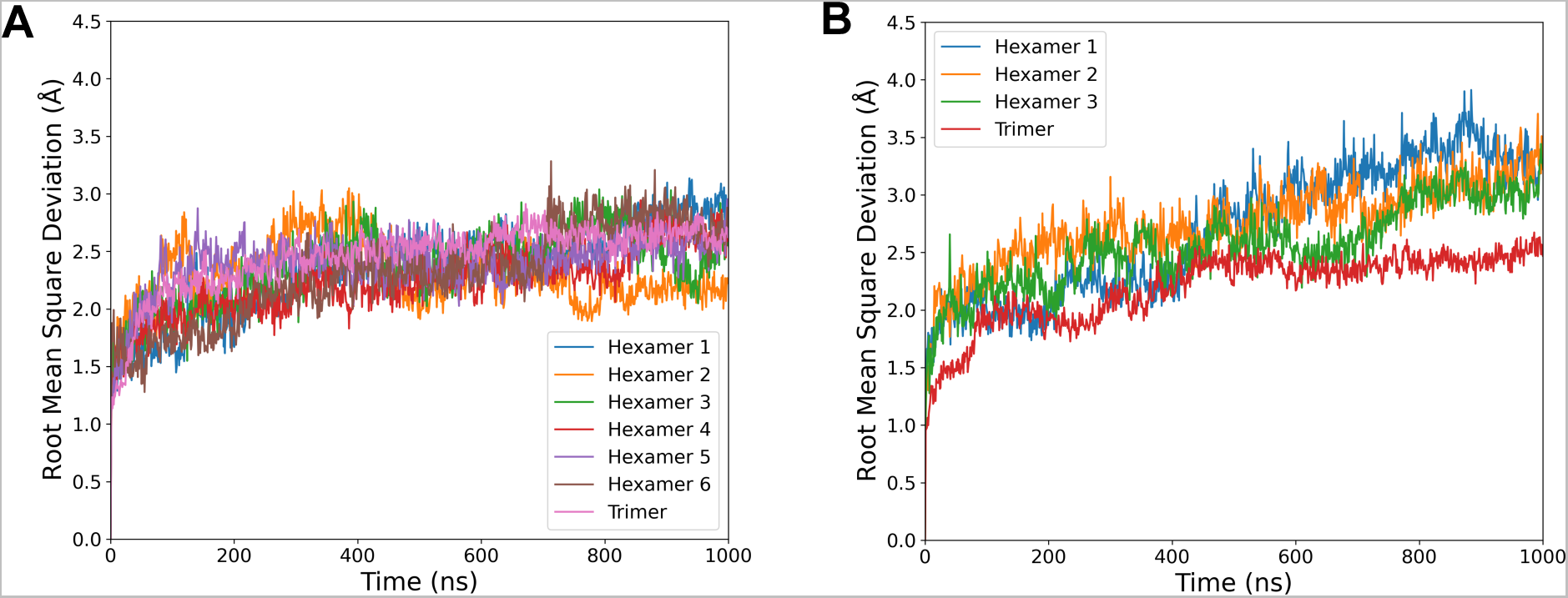
Root mean square deviation (RMSD) of carbon alpha for each hexamers and trimer when taken from the (A) shell fragment or (B) sheet simulation systems from Fig. 3. The reference structure when assessing the RMSD was a tile from initial 6N07 structure,^22^ and is plotted individually for each individual tile within the system.

### Pore Dynamics in BMC Shell Fragments and Sheets

The small variations in RMSD from Fig. 4 leave open the possibility that the individual pores may change their structure when shell proteins are exposed to different local environments. Because pore size and dynamics can alter the metabolite transport, monitoring pore fluctuations over time is of critical importance. On average, we find that the pore radii, both in their ranges and their average, are highly consistent between simulations run in either condition (Fig. 5). On average, the hexamer tile has a central pore with a bottleneck diameter of 6.9 Å in the shell, or 7.1 Å in the sheet. This minimal change indicates that the hexamer pore is invariant to local protein environment, and exhibits similar variation in size across the simulations.

**Figure 5:**
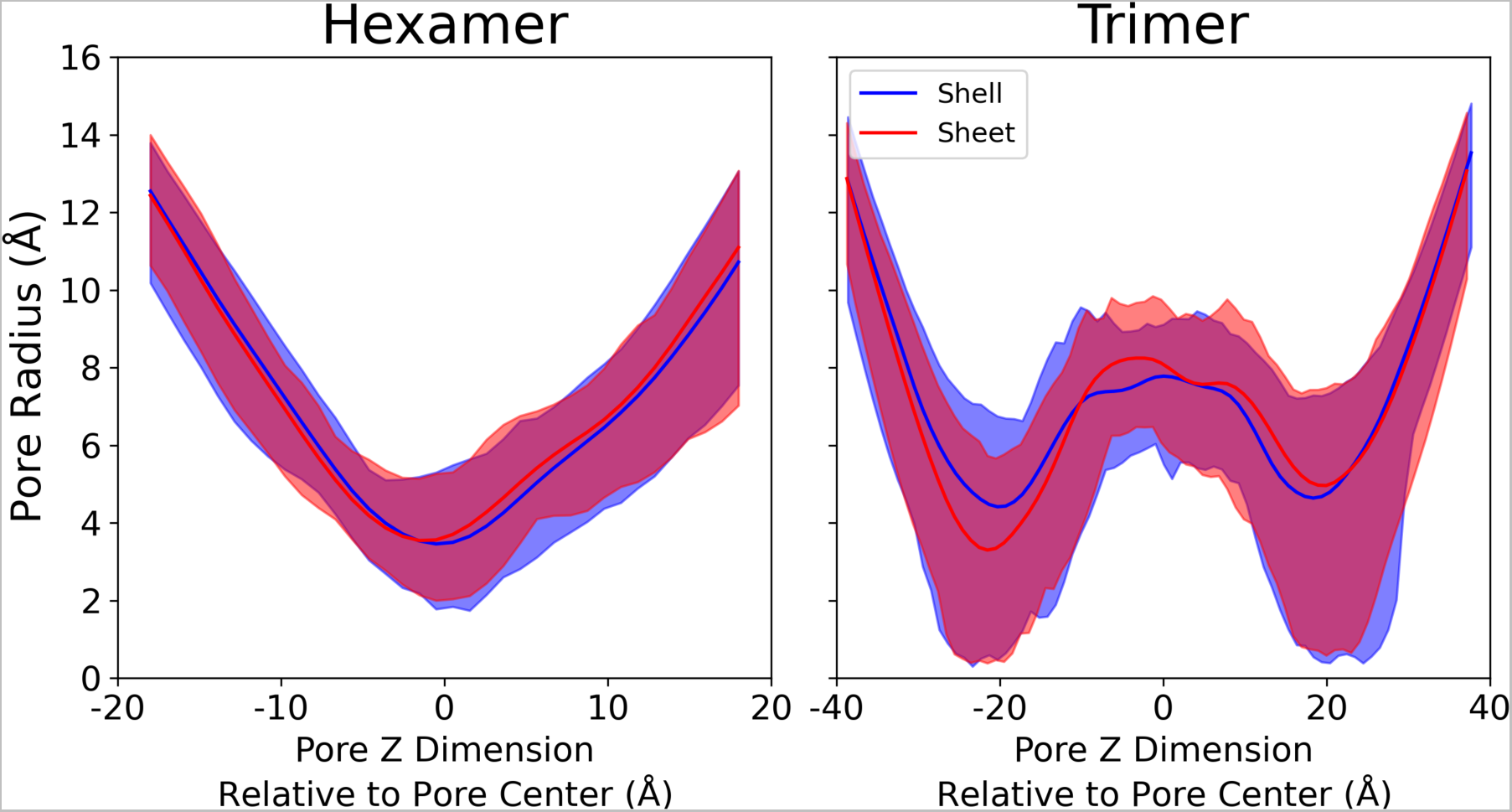
Distribution of pore radii along the central pore for hexameric and trimeric proteins tiles in either a shell-fragment or sheet morphology. In this representation, the midpoint Z is at the geometric center of the hexamers or trimers that make up the pore, which is not where the bottlenecks occur. The shaded area represent the maximum and minimum distribution of pore sizes observed during the simulation. The solid line represent the average pore size observed in the simulation trajectory, across all 6 hexamer tiles (shell fragment) or 3 hexamer tiles (sheet).

The stacked trimer pores exhibit substantially greater variation in the range of possible pore diameters at the bottleneck, at *±*20 Å. In some conformations, particularly at the beginning of the simulation where the structure has not diverged very far from the closed starting point created by the 6N07 starting structure, the trimeric pore is effectively closed at the bottleneck. This closed pore would likely represent a large barrier to permeation to all but the smallest of molecules. However, in other conformations, the trimeric pore has a substantially larger diameter. On average the trimers exposes a larger pore for metabolites to transit across. Thus, we anticipate that the trimer may be the preferred path for some molecules to permeate that cannot be accommodated by the smaller hexamer.

For small molecule permeation, pore dynamics are essential. As the starting structure is taken from a closed starting structure, our initial structures exhibit a closed conformation for the hexamer and trimer complexes. The minimum hexamer opening often occurs at time zero, starting with a bottleneck radius of only 2 Å (SI Animation 1-4). The bottleneck expands quickly as the pore hydrates and sidechains rearrange to a typical radius of 5-6 Å (Fig. 5). The trimeric complex exhibits even stronger dynamics, with an effectively closed pore in the initial structure.^22^ The HOLE output highlights an expanding pore over time (SI Animation 3-4). Once opened, the pore is not observed to dehydrate and close over our 1 µs simulation duration. It is possible that the pore may reclose if the simulation were to be extended, however given how few trimers we have in our simulation systems, the cost in computer time to observe reclosing events was thought to be prohibitive to depend on stochastic sampling to reclose the pore.

### Water Interaction Analysis

Beyond the proteins themselves, our simulation systems feature water to fill in the rest of the simulation volume. While the pore dynamics are likely most critical to permeability, observing changes in water interactions may be another avenue by which we can tease apart the subtleties of structural differences between sheet-like and shell-like structures. The per-residue water contacts vary minimally between the two tested conformations for the hexamer (Figure 6A), suggesting that even structural details are largely conserved at a global scale between the curved shell fragments and a larger sheet. This total picture of conserved contacts is retained if we expand our view to also include the trimer. By mapping water contacts onto the structure, as in Figure S3, we visually see the same water contact patterning across both structures. Contacts are naturally highest on the protein periphery that are solvent exposed, with the residues at the central bottleneck having roughly half the number of water contacts due to protein occluding many potential water interaction sites. In general, this suggests that both structures are facing the same solvent environment, regardless of the exact geometry at play.

**Figure 6:**
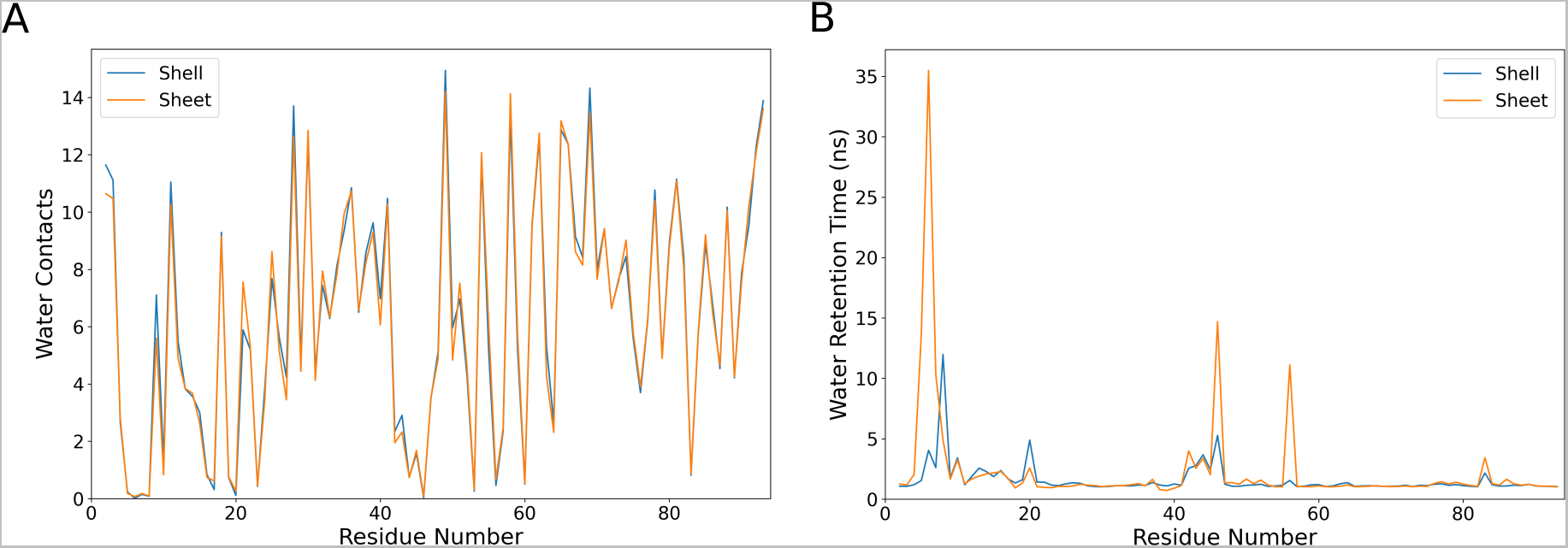
Hexamer tile interactions with water. (A) Quantifies the water contacts on a per-residue basis, counting the average number of pairwise contacts between amino acid residue atoms and water atoms that are under 5 Å in separation over the total trajectory. To maintain consistency with the semi-infinite sheet, the shell values reported here are averaged only over the three hexamer monomers nearest to the central trimer. (B) Measures the water retention times, based on how long on average a given water molecule remains within 5 Å of a given residue. Note that both quantities within the hexamer are averaged over the 18 monomers that are not solvent exposed for the shell fragment system, and over all 18 monomers in the sheet system.

Quantifying contacts alone is only one metric of interest. With an eye towards comparing to hydroxy radical footprinting data, where the timespan spent near specific residues is of interest, we quantify the residence time for water molecules near individual residues (Fig. 6B). We see some increased retention of water molecules near specific residues, with substantial differences in the range of residues 5-11 (LGMIEVR), 20 (A), 41-48 (YVTAVRGD), 50-52 (VAA) and 83 (V) (Figure 6B). The strongest difference in water retention between sheets and shells occurs around G6, where water in the sheet conformation is retained for over 30 ns on average. This residue is on the border of an interstitial water site, near β sheets within an individual hexamer (Figure 7 and 8). The residues lining the pores have much shorter water interactions, as water at the interface readily exchanges with the bulk.

**Figure 7:**
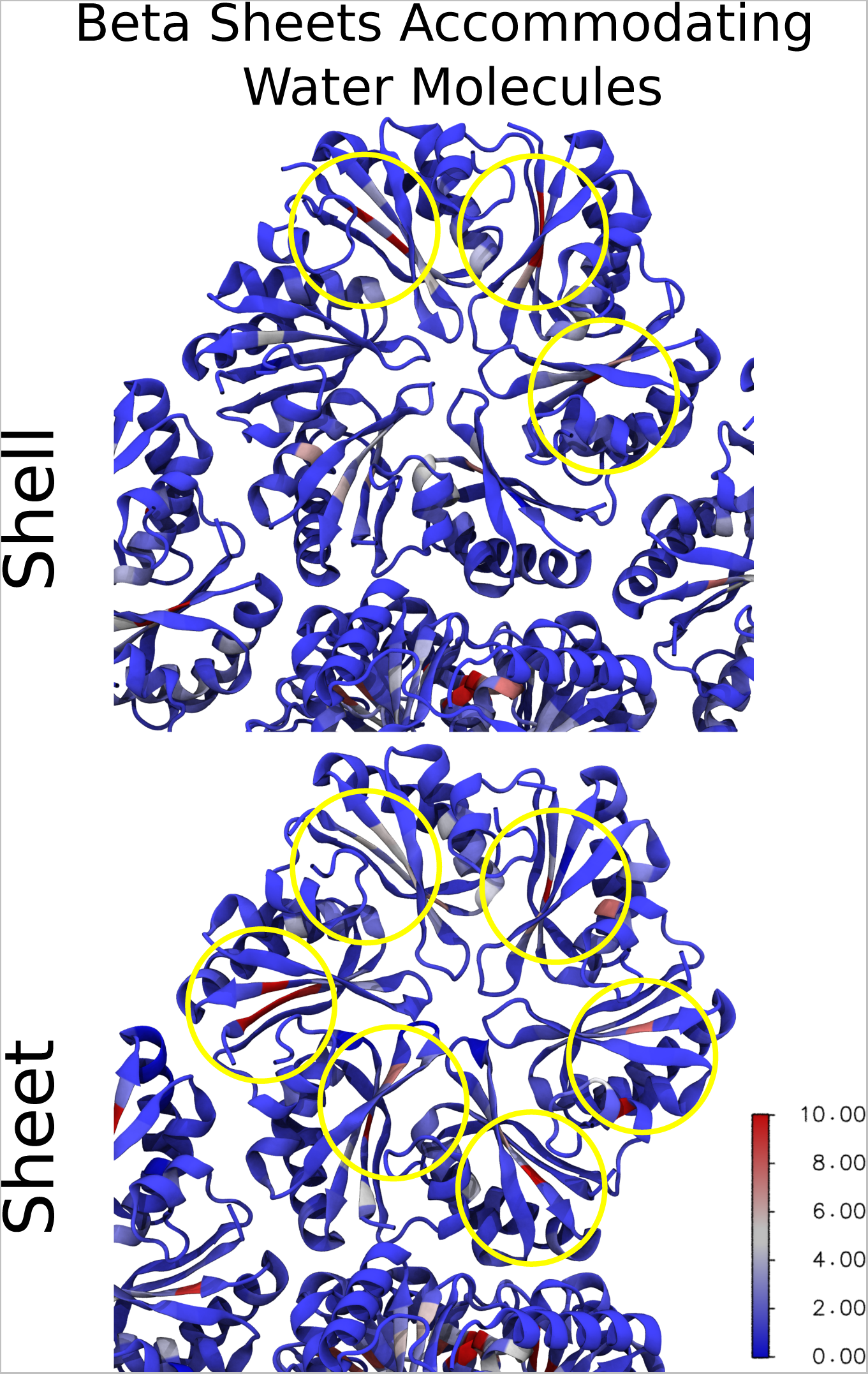
Water retention mapped to each residue on the hexamers and trimers in shell-like and sheet morpholigies. Proteins are drawn in a cartoon representation where each residues is color coded on the blue-white-red spectrum to represent the water retention time. The color bar measures the water retention time in nanoseconds.

**Figure 8:**
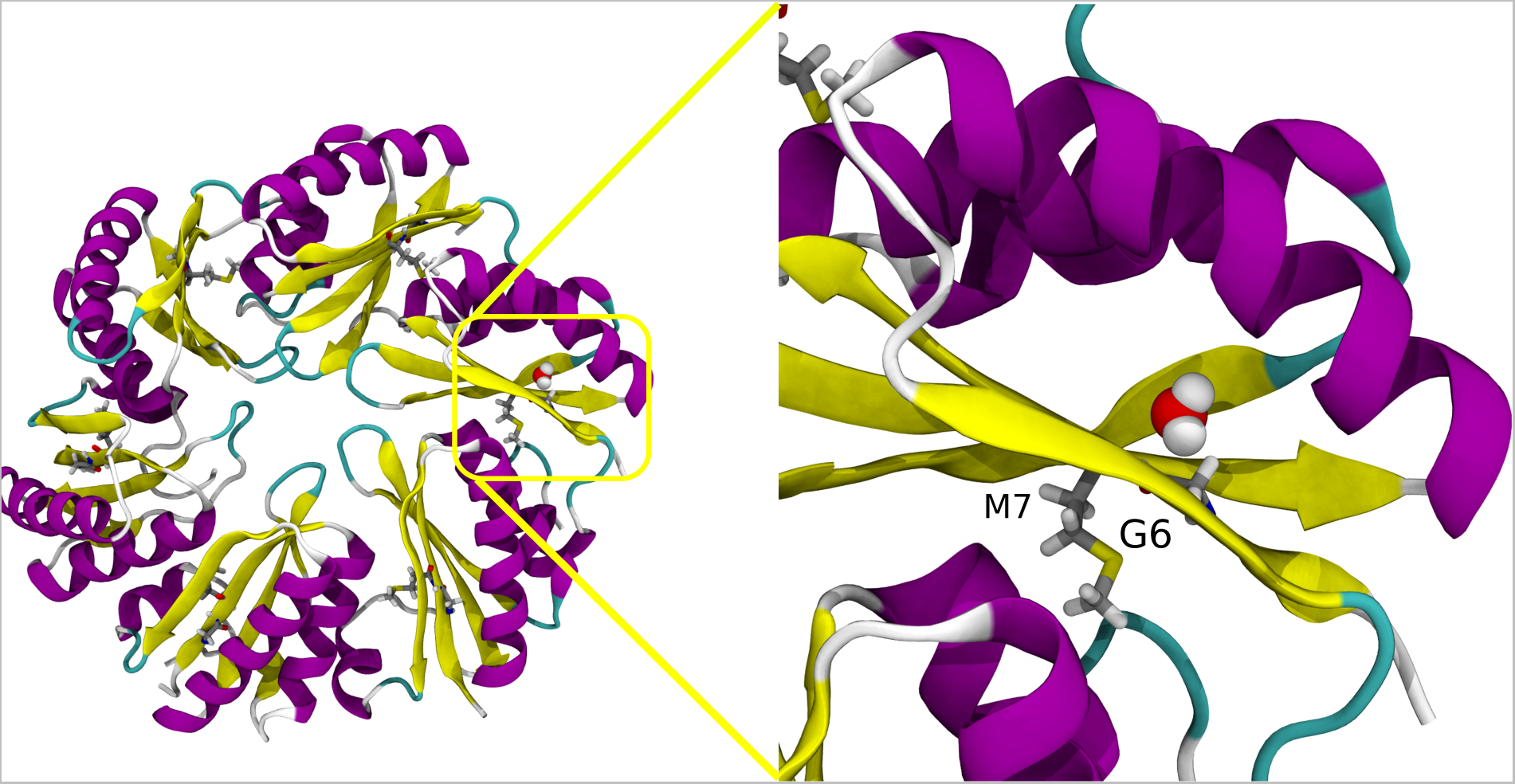
Retention of a water in a pocket around beta sheet in hexamers near residue G6 and M7. Protein are represented in cartoon representation, residue G6 and M7 in licorice and water in VDW. Left panel represent the whole hexamer unit and right panel represents the water pocket.

### Correlating Simulation to XFMS

Our *in silico* work so far has emphasized that the differences between water access and pore formation are really minimal. Is it possible to use an experimental measure like XFMS, where amino acid hydroxylation induced by water ionization through x-ray exposure can be measured by mass spectrometry,^50,51^ to provide experimental support for these findings? Data for the hexamer in solution or as part of a shell is provided in Table 1. The k-values measure the hydroxyl replacement rate at specific residues where hydroxylation is possible. The replacement rate in solution can be greatly accelerated when compared to the complete shell, such as at the M7 residue, which has a very high ratio of hydroxylation rate in solution compared to when it is in the shell. This indicated that this residue has substantially lower availability to water when it is in the shell compared with the solution state. We did not directly simulate an isolated hexameric tile in solution, and not all hexamer monomers within our curved shell fragment are a good facsimile for the environment of a complete shell. Thus, we segment our data into two populations based on hexamer identity (Fig. S2) to establish comparisons with Table 1.

**Table 1:** XFMS k-values at different positions on the BMC hexamer for a single hexamer tile in solution and as part of a BMC shell assembly. The ratio of these two rates tells us something about how water accessibility changes based on BMC protein morphology.

| Residues | Solution k-value | Shell k-value | Ratio |
| --- | --- | --- | --- |
| M7 | $(242.0 \pm 9.6) \times 10^{-6}$ | $(29.8 \pm 8.2) \times 10^{-6}$ | 8.12 |
| M16,23 | $(65.2 \pm 210.0) \times 10^{-6}$ | $(51.2 \pm 1.4) \times 10^{-6}$ | 1.27 |
| Y34 | $(14.7 \pm 1.9) \times 10^{-6}$ | $(155.0 \pm 7.1) \times 10^{-7}$ | 0.95 |
| Y41 | $(6.0 \pm 1.7) \times 10^{-6}$ | $(39.1 \pm 5.5) \times 10^{-7}$ | 1.53 |
| K54 | $(140.0 \pm 8.0) \times 10^{-7}$ | $(64.7 \pm 3.1) \times 10^{-7}$ | 2.16 |
| P77,P79,P88 | $(477.0 \pm 9.1) \times 10^{-6}$ | $(187.0 \pm 8.9) \times 10^{-6}$ | 2.54 |

After this segmentation process, can make comparisons with structural metrics. The Solvent Accessible Surface Area (SASA) for individual residues has been previously demonstrated to correlate somewhat with hydroxyl radical footprinting data derived from XFMS.^52^ There certainly is a trend when evaluating the SASA overall for specific residues (Fig. 9A&B). The fit improves if M7 is excluded from consideration, which might be reasonable as our SASA determination algorithm cannot find a water accessible surface near this buried residue. However, from Figs. 6–7, we know that water can access these buried residues near the β-sheet. Moreover, if water accesses these residues, they may be present near these amino acids for a considerable duration (Fig. 6B). Thus, the initial outlier of M7 may be explained by tracking how long a water molecule is present, as this ratio can change substantially for buried residues (Fig. 9C). Indeed, taking the ratio yields a very strong correlation coefficient (Fig. 9D), although this is almost entirely due to the differential water retention around M7 previously noted (Fig. 6B & 7).

**Figure 9:**
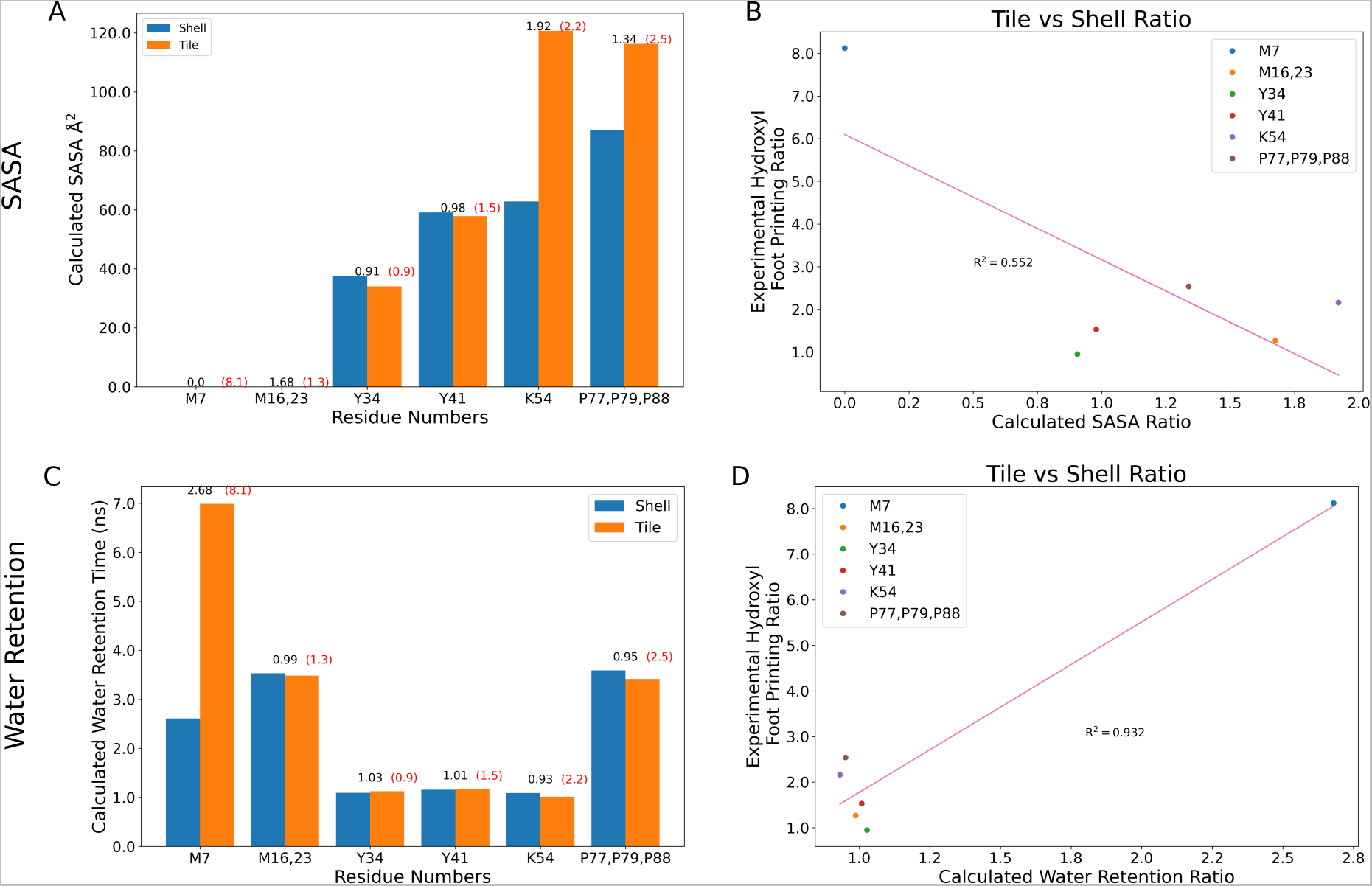
Measured Solvent Accessible Surface Area (SASA) and water retention times across the residues where we have XFMS comparison data (Table 1), directly comparing the equivalent residues within our models that belong to exposed or buried monomers (Fig. S2). The ratios of these quantities determined from simulation are reported in black above the histograms for (A) SASA or (C) water retention times, while the experimental equivalents are written in red. The scatter plot comparing the reaction rate ratios to the ratio of (B) SASA or (D) water retention times have a line of best fit along with a correlation coefficient given in pink.

## Discussion and Conclusions

From the outset, the primary question we were seeking to answer was if future calculations aimed at determining permeability at the molecular scale could assume that the pores are similar irrespective of their local environment within the shell. The direct evidence indicates that this is a reasonable assumption, with Fig. 5 showing little difference in the pore diameter, regardless of whether the hexamer and trimer tiles are arranged as they would in the HO shell, or if they are instead tiled into a planar sheet. Given the substantial reduction in system size that a planar sheet-like arrangement provides, we anticipate using these results as the foundation for the computational simplification of determining permeability through the individual pores at the center of the abundant trimer and hexamer tiles found in many BMC shells.

Indeed, the depth at which we had to look to find any differences between shells and sheets is quite remarkable. The water contacts are the same (Fig. 6A), and while we do not show it, the SASA analysis is also highly similar between sheets and shells when you look at monomers without a solvent-exposed edge. The only difference we find is that there are sporadic trapped waters whose lifetimes are a bit longer in the sheet rather than in the shell (Fig. 6B). Admittedly, the number of trapped water molecules that contribute to these long lifetimes is not large compared to the total number of water molecules that interact with BMC components. However, since we have multiple copies of the hexamer within our system, and can average over 18 monomers where waters may be trapped when conducting our analysis, we are confident that the effect is real.

Our confidence is increased by comparisons to XFMS data for isolated hexamer tiles relative to hexamers within an assembled shell (Table 1). Inferring as we do in Fig. S2 that our shell fragment simulation has components that are in similar environments to both experimental systems, with a solvent exposed edge and a buried edge to individual hexamers, we can readily correlate observed changes in inferred solvent protection from experiment to nanoscale interactions with water (Fig. 9). In particular, the correlation between long-lived water molecules near buried sites such as M7 and the change in the rate of hydroxylation is far stronger than we had anticipated. Initially, the M7 result was thought to be an outlier, but only by using molecular simulation to visualize trapped water molecules can we develop a rational basis for this result.

Zooming out, we think it is helpful to analogize how these BMC protein pores compare with typical membrane channels and transporters. Membrane transporters often must go through a conformational change to fulfill their function.^53^ While we do see pore dynamics over our simulation, with the bottleneck radius increasing and decreasing over time (SI Animation 3-4), we find that these dynamics are primarily driven by sidechain rearrangements, rather than large scale conformational change as might occur in a membrane transporter. Metabolite driven gating has been postulated for other trimer pores,^54–56^ and may well be what occurs in HO shells as well.

Thus, the closest membrane protein analogy for these BMC shell components appears to be that of a channel. Despite starting from a closed state, the trimer opens spontaneously during our simulations, suggesting that the trimer can be gated depending on conditions, analogous to gated ion channels. This was also considered as an explanation for the two particle classes observed cryo-EM studies of the synthetic HO shell. ^22^ The hexameric assembly has a smaller variation in pore diameter, and is more analogous to a constitutively open channel, such as some aquaporins.^57,58^ While the border between channels and transporters is often murky, channels typically have higher conductances.^59^

With the membrane channel analogy in mind, we anticipate that many small molecules may transit through the central pores with high permeability. The limiting factor will be molecular size, as we anticipate that sufficiently large molecules will be unable to transit the pore through these tiled arrangements of BMC shell proteins. Now armed with a computationally efficient planar arrangement of BMC shell components, we are well positioned to test the high permeability hypothesis explicitly. When tested over multiple metabolic pathways featuring different substrates and products, we hope to develop general rules for transport across BMC shells.

## Supporting Information Available

The Supporting Information (SI) contains three figures that due to their size do not fit well into the primary document. These figures show an alternative view of Fig. 3, the segmentation to make the comparison to XFMS data, and mapping the water contacts onto each residue. The SI also has captions for the four animations. All input scripts to build and run molecular simulations are made publicly available on Zenodo. ^26^

## Supporting information

Animation S1

Animation S2

Animation S3

Animation S4

## Acknowledgement

We thank Jonathan K. Lassia for providing samples for the XFMS work. Research by R.S., J.M., M.S., C.A.K., C.Y.R. and J.V.V. was supported as part of the Center for Catalysis in Biomimetic Confinement, an Energy Frontier Research Center funded by the U.S. Department of Energy (DOE), Office of Science, Basic Energy Sciences (BES), under award DE-SC0023395. The Advanced Light Source and the Molecular Foundry are supported by the Office of Science of the U.S. Department of Energy (DOE) under contract DE-AC02-05CH11231. XFMS work was supported by NIH R01 GM126218 and NIH P30 GM124169. D.S. is supported by the U.S. Department of Energy, Office of Basic Energy Sciences under grant number DE-FG02-91ER20021. This research used resources of the National Energy Research Scientific Computing Center (NERSC), a U.S. Department of Energy Office of Science User Facility located at Lawrence Berkeley National Laboratory, operated under Contract No. DE-AC02-05CH11231 using NERSC award BES-ERCAP0024035. Preliminary simulations were supported in part through computational resources and services provided by the Institute for Cyber-Enabled Research at Michigan State University.

## Notes

https://zenodo.org/doi/10.5281/zenodo.10800569

### Competing Interest Statement

The authors have declared no competing interest.

