## Supporting information pdf for "Comparative Pore Structure and Dynamics for Bacterial Microcompartment Shell Protein Assemblies in Sheets or Shells"

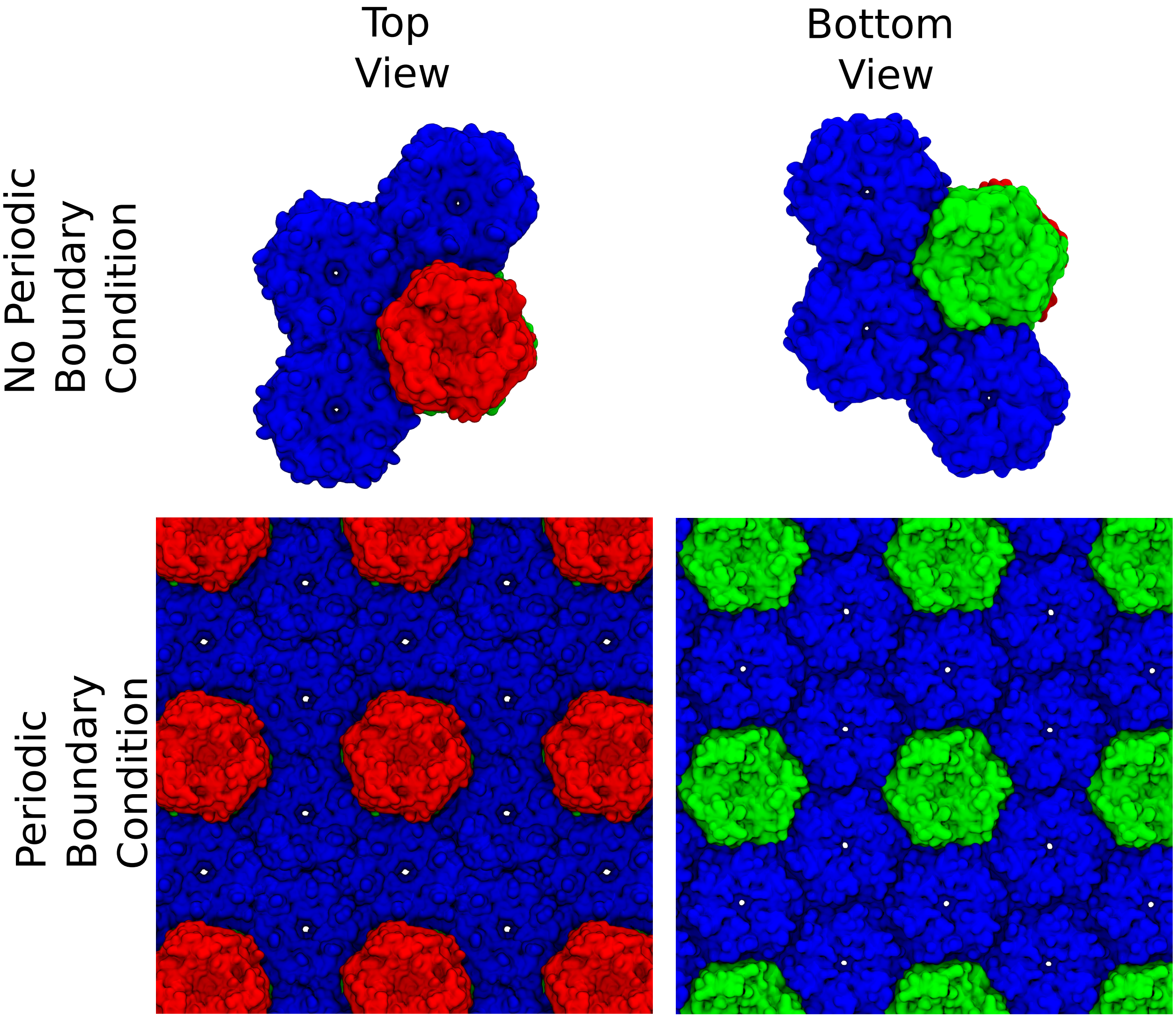


Figure S1: Reduced simulation system consisting of three hexameric units and one trimer dimer which can mimic the planar sheet conformation and keeps the system periodic in a conventional orthogonal unit cell. Hexamers are shown in a blue surface representation, while the trimer upper half and lower half are red and green surfaces, respectively.

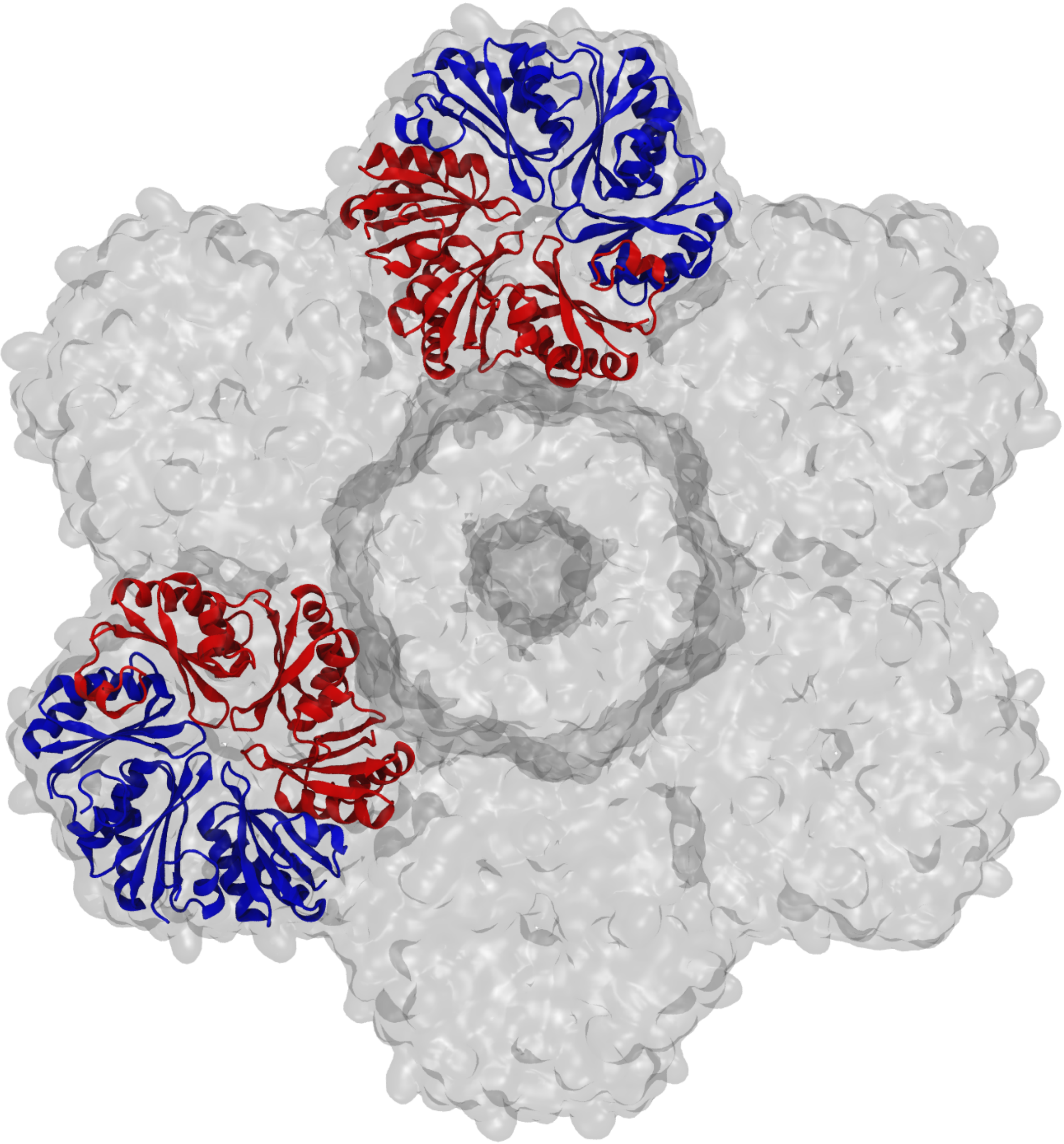

Figure S2: Exposed (blue) and buried (red) hexamer monomers from our shell-fragment simulation. The exposed hexamers and buried hexamers are quantified independently, with the exposed hexamer monomers standing in for a well solvated hexamer tile and the buried hexamers are used to determine averages for a complete shell.

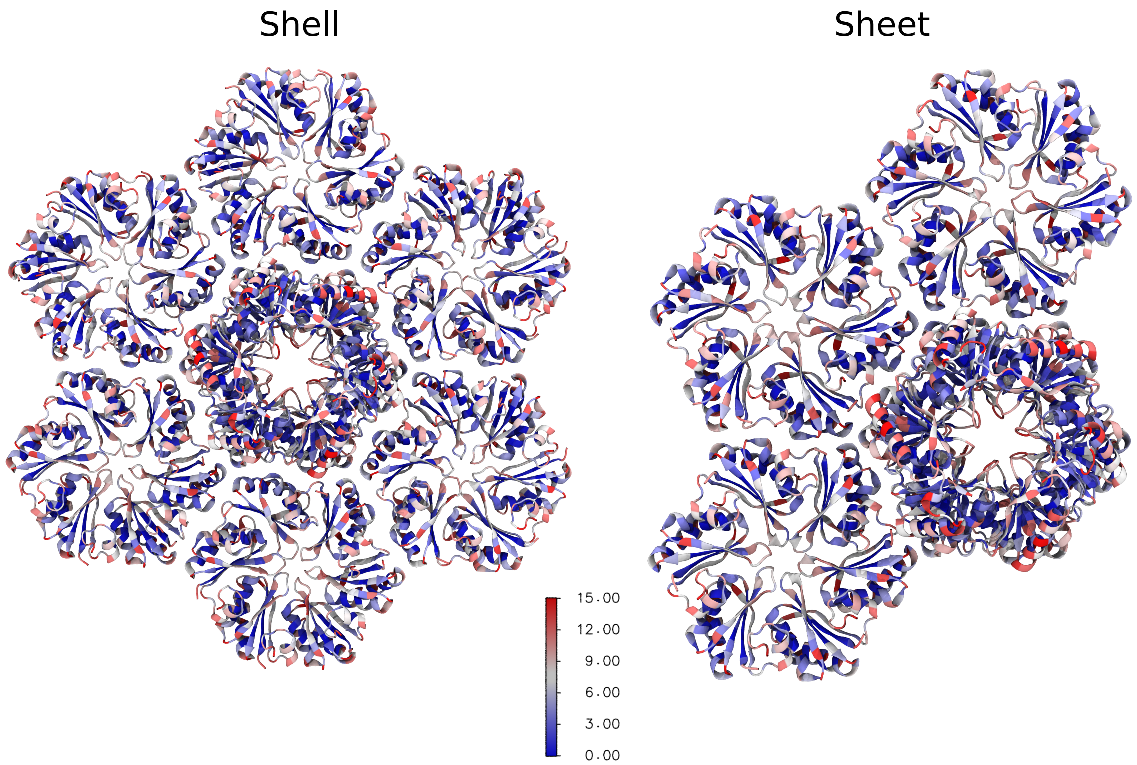


Figure S3: Water contacts mapped to each residue of the shell fragment and sheet assemblies. The protein is represented as a cartoon and color coded on the blue-white-red spectrum with blue representing no contacts and red 15 or more contact number with water.

### SI Animation Captions

1. **Supplementary Animation S1**: Molecular dynamics simulation for the sheet system featuring hexameric and trimeric BMC shell tiles. To make clear that the structure has no gaps, this representation includes periodic images. The animation uses the same representation as Fig. S1, and is a top view to look down all the pores simultaneously.

2. **Supplementary Animation S2**: Molecular dynamics simulation of shell fragment from a larger BMC. The animation uses the same representation as Fig. S1, and is viewed from the top of the shell fragment to facilitate looking down along the trimer pore. Note that due to the curvature of the shell fragment, you cannot see clearly down the hexamer pores.

3. **Supplementary Animation S3**: Animation depicting pore dynamics in BMC trimer (top) and hexamer (bottom) pores when taken from a shell fragment. The left panel highlights the pore as calculated by HOLE for that particular snapshot over time. The right panel quantifies the pore radius along the Z dimension relative to the protein center. Note that while only one hexamer pore is shown in this animation, multiple pores contributed to the analysis in Figure 5.

4. **Supplementary Animation S4**: Animation depicting pore dynamics in BMC trimer (top) and hexamer (bottom) pores when taken from a planar shell sheet. The left panel highlights the pore as calculated by HOLE for that particular snapshot over time. The right panel quantifies the pore radius along the Z dimension relative to the protein center. Note that while only one hexamer pore is shown in this animation, multiple pores contributed to the analysis in Figure 5.
